# Preclinical evaluation of Brincidofovir in glioblastoma demonstrates improved long term-survival and cytomegalovirus-dependent and independent effects

**DOI:** 10.64898/2026.08.20.746020

**Authors:** Noe B. Mercado, Philippa Vaughn-Beaucaire, William M. Hawkins, Andrea Schmidt, Jasmine Clark, Michelle Shub, Mariia Vorobeva, Yovany Padilla, Austin Jacobson, Ayaan Akhtar, Paola Sundaram, Eleni Panagioti, Eain A. Murphy, James A. Lederer, Masatoshi Hazama, Charles H. Cook, Sean E. Lawler

## Abstract

Cytomegalovirus (CMV) has been implicated in glioblastoma (GBM) progression. Ongoing clinical trials are assessing therapeutic approaches targeting CMV in GBM but to date no new therapy has been approved outside the standard of care. Previous preclinical studies have highlighted the potential of the antiviral drug Cidofovir (CDV) in GBM; however, its clinical use is limited by dose-dependent nephrotoxicity and poor cellular uptake, necessitating high intravenous doses to achieve therapeutic activity. Brincidofovir (BCV), a lipid conjugate of CDV has been developed, which does not induce nephrotoxicity and has significantly greater cellular bioavailability. Here we examined the effects of BCV in a newly established CMV-driven GBM model (SB28) and in patient-derived tumor neurospheres. We show that BCV prolongs survival *in vivo* and exerts both CMV-dependent and independent antitumor effects. Mechanistically, BCV induces DNA damage and cell cycle dysregulation in GBM cells and inhibits proliferation of patient-derived neurospheres in a dose-dependent manner. These data identify BCV as a dual-action therapeutic that suppresses viral oncomodulation while directly targeting tumor cell viability.

## INTRODUCTION

Glioblastoma (GBM) is a deadly brain tumor with therapies limited in their efficacy^1–3^. The current standard of care (SOC) treatment for GBM involves surgical resection followed by radiotherapy and chemotherapy using the alkylating agent temozolomide (TMZ)^4^. However, GBM patients face a grim prognosis with only a 15-month median survival after diagnosis. The tumor microenvironment (TME) of GBM is highly heterogenous in cellular and molecular composition. Of importance for immunotherapy approaches is that the TME is heavily infiltrated by immunosuppressive myeloid derived cells including macrophages often referred to as M2-like tumor-associated macrophages (TAMs)^5^. These TAMs promote immunosuppression, invasion, and angiogenesis through the production of anti-inflammatory cytokines. Conversely, other immune cells such as CD8^+^ T cells and CD4^+^ T cells are scarce in the TME^6^. The compositional heterogeneity of the GBM TME makes therapeutic drug development difficult and to date unsuccessful.

Recent studies indicate that targeting Human Cytomegalovirus (HCMV), a widespread DNA virus, in GBM patients may be a promising approach^7–9^. HCMV is a ubiquitous β-herpesvirus that establishes lifelong latency after primary infection^10^. Latent infection of CMV is established in bone marrow hematopoietic progenitor cells with reports suggesting endothelial cells may be another reservoir^11,12^. Recent reports have found HCMV proteins and nucleic acids in GBM specimens^13–16^. In addition, HCMV seropositivity has been correlated to poor survival in GBM patients and has also been shown to promote stemness, invasion, and proliferation^17–19^. Accumulating clinical data support the relevance of CMV in GBM^20–22^ with HCMV-targeted therapeutic approaches underway. A recent clinical trial of newly-diagnosed GBM patients treated with the antiviral drug Valganciclovir (VGCV) with SOC resulted in a significant improvement in overall survival^23^. A larger follow-up clinical trial (NCT04116411) is currently recruiting patients. However, there is limited data on whether other antiviral drugs may therapeutically target GBM *in vivo* and importantly how CMV drives GBM tumor progression and whether it can be therapeutically targeted by these antiviral drugs.

Mouse models of GBM can provide an important platform to investigate how CMV contributes to tumor progression and evaluate therapeutic strategies targeting viral oncomodulation. Previously, we reported that MCMV infection in the GL261 murine GBM model promoted tumor progression and angiogenesis, resulting in a significantly reduced survival compared to uninfected controls^24^. However, the GL261 model does not fully recapitulate human GBM because of its high mutational burden and relatively immunogenic microenvironment. Therefore, here we used the SB28 murine GBM model, which more closely mimics key features of human GBM, including aggressive growth, low mutational burden, and a profoundly immunosuppressive TME^25–27^.

In our previous report we showed that treatment with CDV improved survival in infected mice and inhibited MCMV reactivation as well as tumor angiogenesis^24^. Other studies reported that CDV has antineoplastic activity against HCMV infected GBM cells and treatment with CDV in combination with radiotherapy extended survival in a xenograft GBM model^28^. However, CDV is a poor therapeutic drug due to its dose limiting nephrotoxicity impacting proximal renal tubules^29^. Therefore, administration of CDV requires routine prophylactic measures for safe use to avoid kidney injury. CDV primarily uses the cellular transporter human organic anion transporter 1 (hOAT1) which is expressed on renal proximal tubule cells and results in high intracellular accumulation in the kidney^30^. Conversely, hOAT1 expression is low or completely absent in cells like fibroblast that are readily infected by viruses like CMV. This results in slow and inefficient uptake of CDV where it is therapeutically needed^30^.

To overcome the limitations of CDV, Brincidofovir (BCV), a lipid-conjugate of CDV with improved cellular bioavailability, was developed^31^. BCV delivers higher intracellular concentrations of cidofovir-diphosphate (CDV-PP), avoids the nephrotoxicity associated with CDV, and exhibits broad antiviral activity against double stranded DNA viruses, including CMV. CDV-PP structurally resembles dCTP, a substrate of viral DNA polymerase and can be incorporated into the viral genome thereby resulting in inhibition of viral DNA replication^18,32^. A Phase I clinical study assessed BCV pharmacokinetics and safety demonstrating that BCV can be safely administered to healthy patients without significant adverse events^33^. Another phase II trial (NTC00942305) showed that treatment with BCV in allogenic hematopoietic-cell transplant (HCT) patients reduced plasma HCMV DNA levels compared to placebo controls^34^. Notably, BCV received FDA approval in 2021 for the treatment of smallpox, underscoring its translational and clinical potantial^35–37^.

In this study, we assessed whether BCV inhibits GBM growth in both the presence and absence of CMV. Using the SB28 murine GBM model, we demonstrated that CMV promotes tumor progression and that BCV significantly prolongs survival in both CMV-infected and uninfected tumor bearing mice. Mechanistically, BCV directly inhibited GBM cell growth in vitro and induced tumor cell DNA damage, supporting both CMV-dependent and independent anti-tumor effects. Collectively, these data identify BCV as a potential therapeutic strategy for GBM and support its further evaluation in future clinical studies.

## RESULTS

### Cytomegalovirus infection accelerates GBM tumor growth in mice

SB28 is a murine model of GBM that closely mimics aggressive human GBM growth and tumor immune suppression. To investigate whether CMV accelerates tumor growth in the SB28 model of GBM, we perinatally infected C57BL/6 mice at P2 with Δm157 MCMV then allowed to resolve for 14 weeks (Figure 1A) as we have previously described^24^. SB28 cells were then intracranially implanted into MCMV infected and age-matched naïve control mice at 14 weeks of age. In this model we observed a significant decrease in survival of CMV infected tumor bearing mice compared to naïve controls (median survival 23 days vs 31 days in naïve, *p* < 0.02, log-rank test) (Figure 1C) along with early onset of GBM clinical signs in the MCMV infected cohort compared to the naïve cohort. Mice infected with MCMV had higher weight loss compared to naïve cohort (Figure 1B). This suggests that MCMV mice experience faster clinical decline compared to naïve mice. Additionally, the MCMV infected cohorts had significantly brighter and larger areas of GFP expression after IVIS imaging analysis suggesting that these tumors are larger compared to the naïve cohort (Figure 1D). At endpoints whole brains were cryosectioned for immunostaining and IVIS imaging for GFP signal. Immunostaining of brain sections for the endothelial marker CD31 revealed a significantly higher number of CD31-positive cells in MCMV-infected tumor cohorts compared to naïve (uninfected) controls (Figure 1E, G). This observation suggests enhanced vascular formation within the MCMV-infected tumors. Notably, CD31 immunoreactivity was predominant in the tumor periphery (edge) compared to the core, which may indicate preferential vascularization. Concurrent staining for MCMV using an anti-m123 (IE1 homolog) antibody revealed detectable viral immediate early protein expression within the infected tumors, confirming MCMV presence. (Figure 1F, H). Thus, consistent with our previous findings in the GL261 GBM model, CMV promotes tumor progression in the clinically relevant SB28 model and is associated with viral reactivation within the tumor microenvironment.

**Figure 1:**
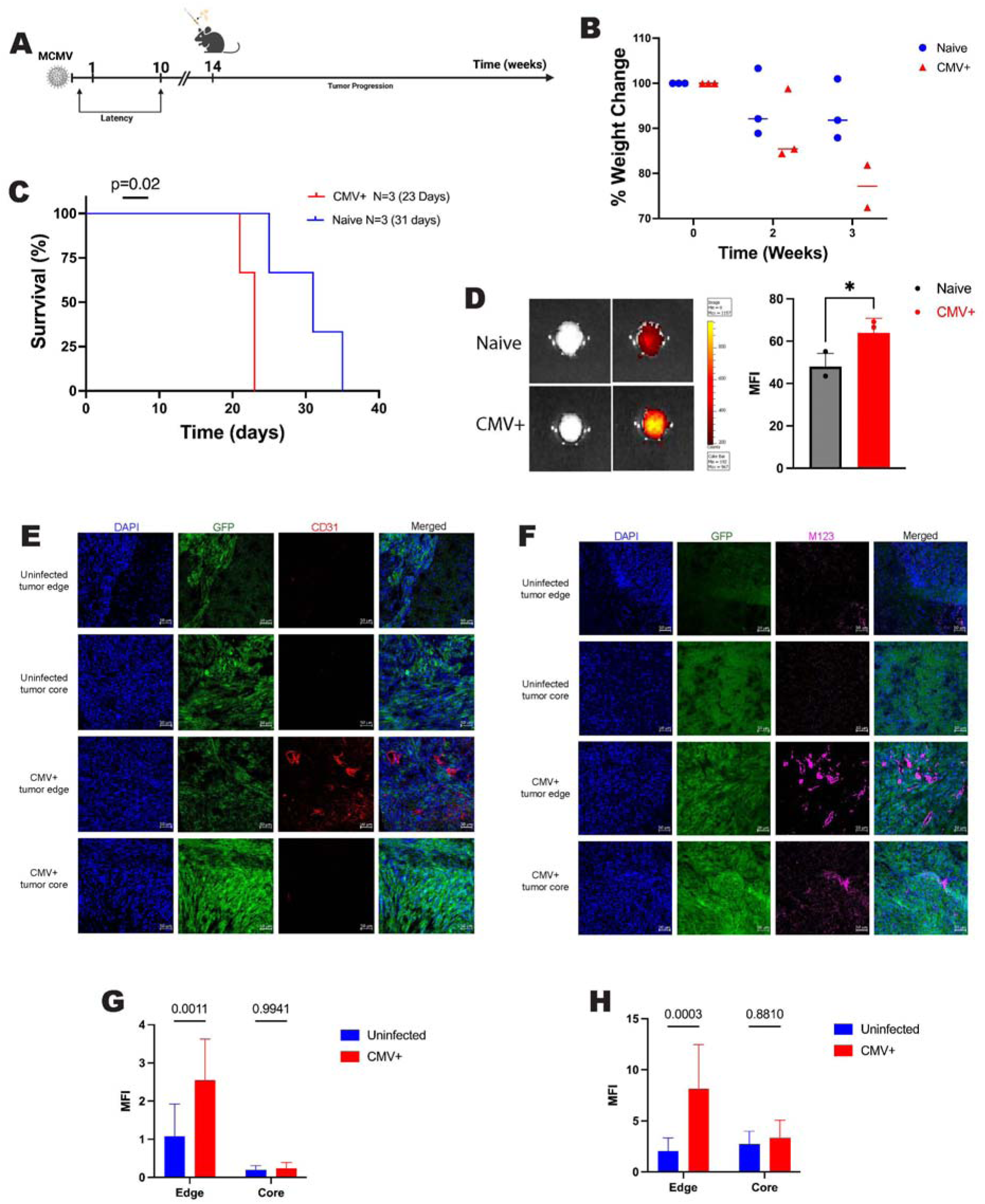
MCMV infection promotes tumor growth and clinical decline in a syngeneic murine model of GBM. (A) Experimental overview of MCMV infection and SB28 tumor implantation. (B) Percent weight loss of naïve-uninfected mice compared to MCMV infected mice after tumor implantation (weekly measurements) (C) Kaplan-Meier curves of SB28 tumor-bearing mice. Naïve-uninfected, *n* = 3; MCMV^+^, *n* = 3. *p*<0.02, log-rank test. Median survival is indicated on plot and shown in parentheses. (D) Representative ex vivo fluorescence imaging of dissected brain from tumor bearing mice using IVIS. Color scale represents fluorescent radiance (photons/s/cm^2^). Scale bar indicates GFP signal intensity as a measure of tumor burden (CMV+ and naïve groups were compared using an unpaired two-tailed Welch’s t-test. Data are shown as individual values with mean ± SD. Statistical significance: * p < 0.05; ns, not significant (p > 0.05). (E) CD31 (red) and (F) MCMV (pink, anti-m123) immunofluorescence of tumor sections from naïve uninfected and MCMV infected mice at end points Scale bar: 50μm. Quantification of (G) CD31 and (F) MCMV M123 mean fluorescence intensity (MFI) in SB28 tumor tissue (two-way ANOVA, Tukey’s multiple comparisons test, mean ± SD. Statistical significance: *, *p* < 0.05; p > 0.05, ns, not significant. n=7 independent fields).

### Brincidofovir prevents cytomegalovirus induced tumor progression and improves survival

Next, we asked whether BCV exerts direct anti-tumor activity in the SB28 murine GBM model. To test this, 1,000 SB28 cells were intracranially implanted in C57BL/6 mice. On day 8 post implantation we initiated BCV treatment 20mg/kg twice weekly for 1 month (8 total doses) then monitored mice for clinical decline. Bioluminescence imaging (BLI) 2 weeks after tumor implantation revealed that 4 out of 5 mice (80%) had positive signal of tumor presence as demonstrated by luminescence intensity (Supplemental Figure 1A). Comparatively only 1 out of 5 mice (20%) in the BCV treated cohort had a luminescence signal, suggesting that tumors in the majority of control animals were below the threshold of detection. We observed a significant increase in the survival of the BCV treated mice compared to the untreated controls (Supplemental Figure 1B) *(p* < 0.0057, log-rank test). Importantly, 20% of BCV-treated mice achieved long-term survival compared to 0% in untreated controls (Supplemental Figure 1). Thus, our data show that BCV has antitumor activity *in vivo* in the SB28 murine model of GBM.

Next, we investigated whether CMV promotes tumor progression and reduces survival and whether this effect can be reversed by BCV treatment. To address this, we perinatally infected C57BL/6 pups at P2 with MCMV (Δm157) then allowed to resolve for <10 weeks (Figure 2A). At week 14 we intracranially implanted 1,500 SB28 cells (Figure 2A). Mice were separated into cohorts, BCV treated and untreated (BCV+/-) and CMV+/-. On day 8 post-tumor implantation, BCV treatment was initiated 20mg/kg twice weekly for 1 month. Mice received a total of 8 doses and no adverse events were observed. We found that MCMV significantly decreased survival of tumor bearing mice (median survival 24 vs 28.5 days in uninfected controls; p<0.0264, log rank test; Figure 2B), and that this effect was reversed by BCV treatment (46 vs 24 days; p<0.0001). Notably, BCV also improved survival in uninfected-tumor bearing mice (49 vs 28.5 days; p<0.0001; Figure 2B) indicating activity independent of viral infection. BCV promoted long-term survival (>80 days) in both MCMV+ (44.4%) and uninfected cohorts (25%). Collectively, these data support dual antiviral and direct antitumor effects of BCV in this model.

**Figure 2:**
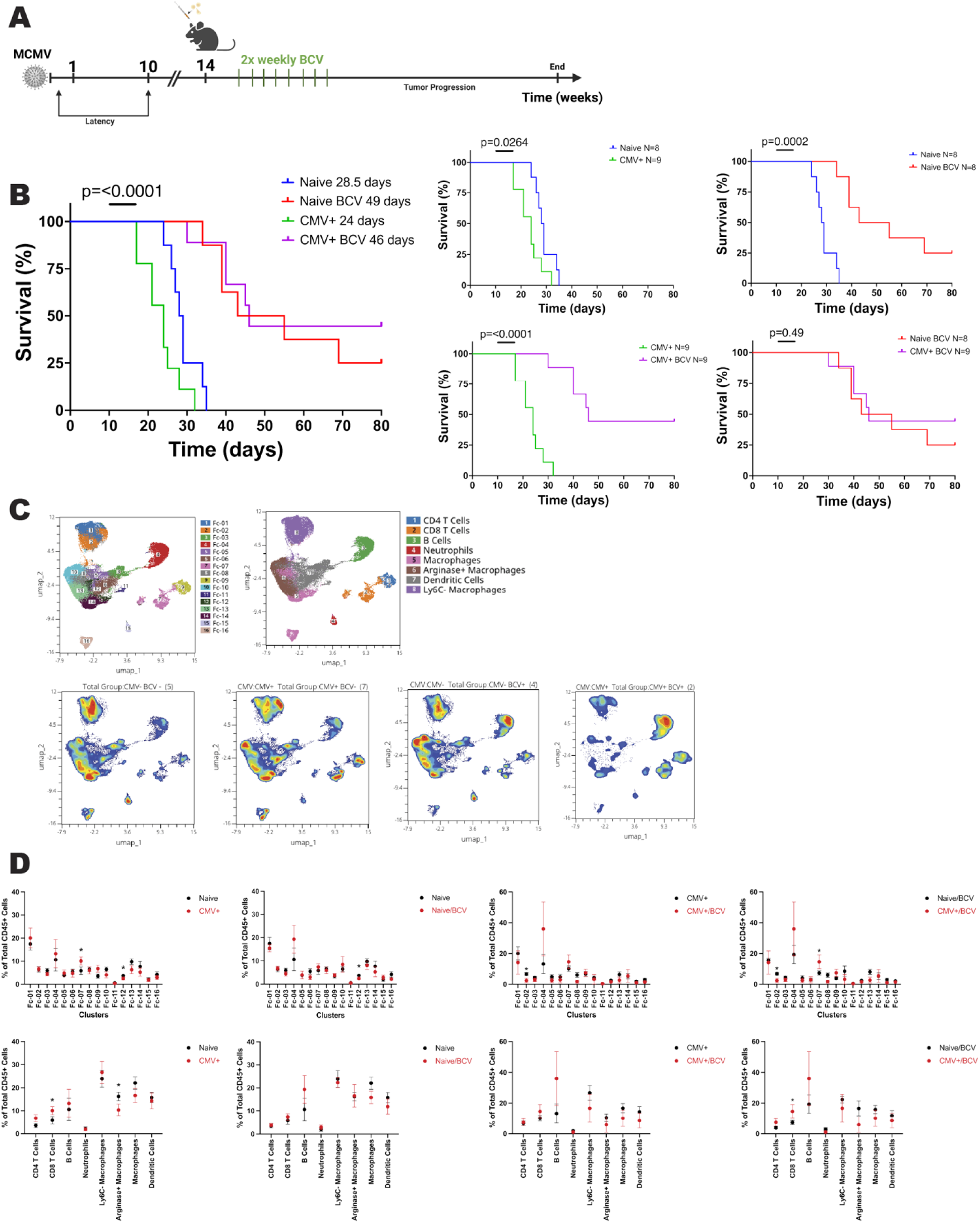
Brincidofovir treatment improves survival of SB28 tumor bearing C57BL/6 mice. (A) Experimental overview. (B) Kaplan-Meier survival curve of naive and MCMV infected, SB28 tumor-bearing mice, four cohorts are compared 1. Naïve-untreated 2. MCMV infected 3. Naïve-Brincidofovir (BCV) treated and 4. MCMV infected Brincidofovir (BCV) treated, *P* < 0.0001, log-rank test. Median survival is indicated on plot and shown in parentheses. Extrapolated plots of the four cohorts, 1. naïve-uninfected vs MCMV infected p<0.0264, log rank test 2. Naïve-uninfected vs Naïve-uninfected treated with BCV p<0.0002, log rank test 3. MCMV infected vs MCMV infected BCV treated p<0.0001, log rank test 4. Naïve-uninfected treated with BCV vs MCMV infected treated with BCV p<0.49, log rank test (C) FlowSOM clustering of CyTOF data identifies 16 distinct immune cell clusters. Single cell suspensions from tumor bearing hemisphere from each of the cohorts were analyzed, Naïve-untreated n=4, MCMV infected n=7, Naïve-BCV treated n=4, and MCMV infected Brincidofovir (BCV) treated n=2. Individual UMAP projections of cells belonging to each of the 16 clusters (one panel per cluster), colored by expression of key phenotypic markers. (D) Plots of clusters and major immune cells subsets (CD4+ T cells, CD8+ T cells, B cells, neutrophils, macrophages (Ly6c+ and arginase+) and dendritic cells. Statistical analysis for Cytof was performed using Student’s T-test. Statistical significance: * *p* < 0.05; p > 0.05, ns, not significant.

### CMV promotes immune remodeling and is reversed by Brincidofovir

To investigate how CMV and BCV modulate the TME, we performed CyTOF analysis of tumor-infiltrating lymphocytes at endpoint. Sixteen cell clusters with distinct marker expression were identified and consolidated into eight major immune cell subsets (Figure 2C). CMV infection was associated with a higher proportion of CD8^+^ T cells and a decreased frequency of arginase expressing macrophages (*p* ≤ 0.05), indicative of a more inflamed TME (Figure 2D). Interestingly, these differences were abrogated by BCV treatment, which instead showed a trend toward increased B cells (p = 0.07), suggestive of enhanced immune surveillance. Direct comparison of BCV-treated cohorts (CMV+ vs CMV-) revealed higher CD8^+^ T cell frequencies in CMV+ mice (*p* ≤ 0.05), supporting a CMV-driven inflammation state. Cluster-level analysis further showed a reduction of Fc-12 dendritic cells in BCV-treated uninfected mice and a decrease in Fc-02 Ly6C macrophages in CMV-infected treated mice (*p* ≤ 0.05). Together, these data suggest that CMV shapes the immune landscape of the TME, while BCV counteracts these effects and restores immune balance.

### Brincidofovir induces S-phase cell cycle arrest and promotes DNA damage in GBM

To mechanistically assess the impact of BCV on tumor cells following CMV infection, SB28 cells and patient derived GBM neurospheres (G44) were infected and subsequently treated with BCV. Cell cycle distribution was analyzed by propidium iodide and flow cytometry. BCV treatment significantly increased the proportion of cells S-phase in both SB28 cells and G44 neurospheres compared to controls (*p* < 0.05; Figure 3A, B). CMV infection of SB28 cells led to increased fraction of cells in G2/M phase and a reduction in G1, consistent with enhanced proliferation (Figure 3A), an effect not observed in G44 neurospheres, potentially reflecting differences in infection kinetics. Indeed, CMV-infected neurospheres showed increased size at day 7 compared to controls (Figure 2B and Figure 4A). Taken together, these data demonstrate that BCV alters GBM cell cycle dynamics.

**Figure 3:**
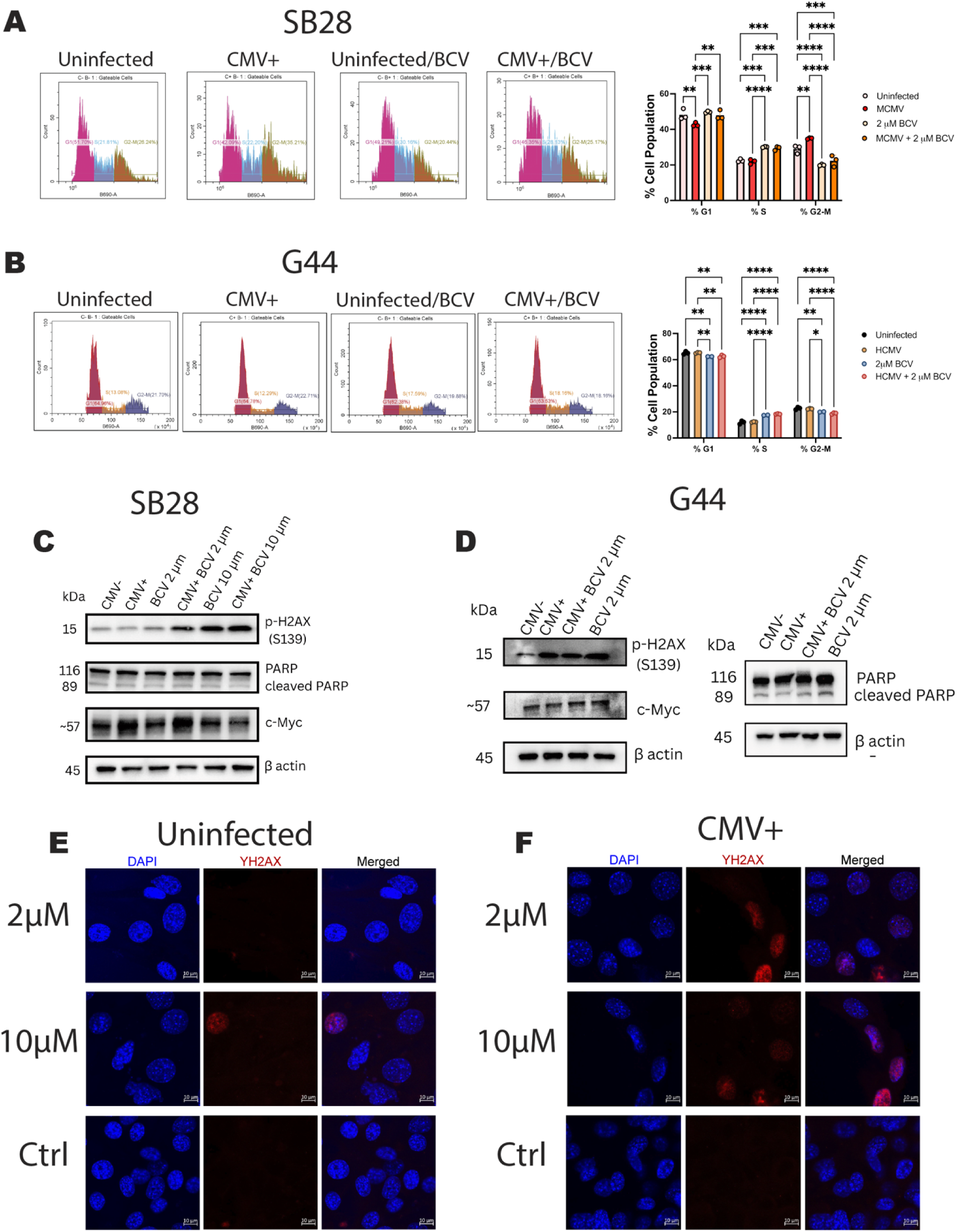
BCV induces cell cycle arrest and DNA damage in GBM. (A) Flow cytometry cell cycle analysis using propidium iodide (PI) DNA staining of tumor neurospheres (G44) and (B) SB28 murine GBM cells. Representative histograms (PI fluorescence intensity (B690-A) on x-axis and cell count on y-axis) comparing naïve-uninfected vs MCMV infected controls and BCV treated. Sequential gating on forward scatter (FSC) vs. side scatter (SSC) to select total intact cells, followed by FSC-height vs. FSC-area for singlet selection. Quantified percentage of cells in each cell cycle phase across control and BCV treated groups. Statistical analysis was performed using two-way ANOVA followed by Tukey’s multiple comparisons test. Data are presented as mean ± SD. Statistical significance: * *p* < 0.05; p > 0.05, ns, not significant. Immunoblot analysis of (C) SB28 and (D) G44 tumor neurospheres, showing levels of phosphorylated γ-H2AX, c-Myc and PARP/cleaved PARP after infection and treatment with BCV. Representative foci of γ-H2AX immunofluorescence staining (red) in (E) SB28 cells and (F) SB28 cells infected with MCMV with nuclei counterstained with DAPI (blue). Scale bar 10μM.

**Figure 4:**
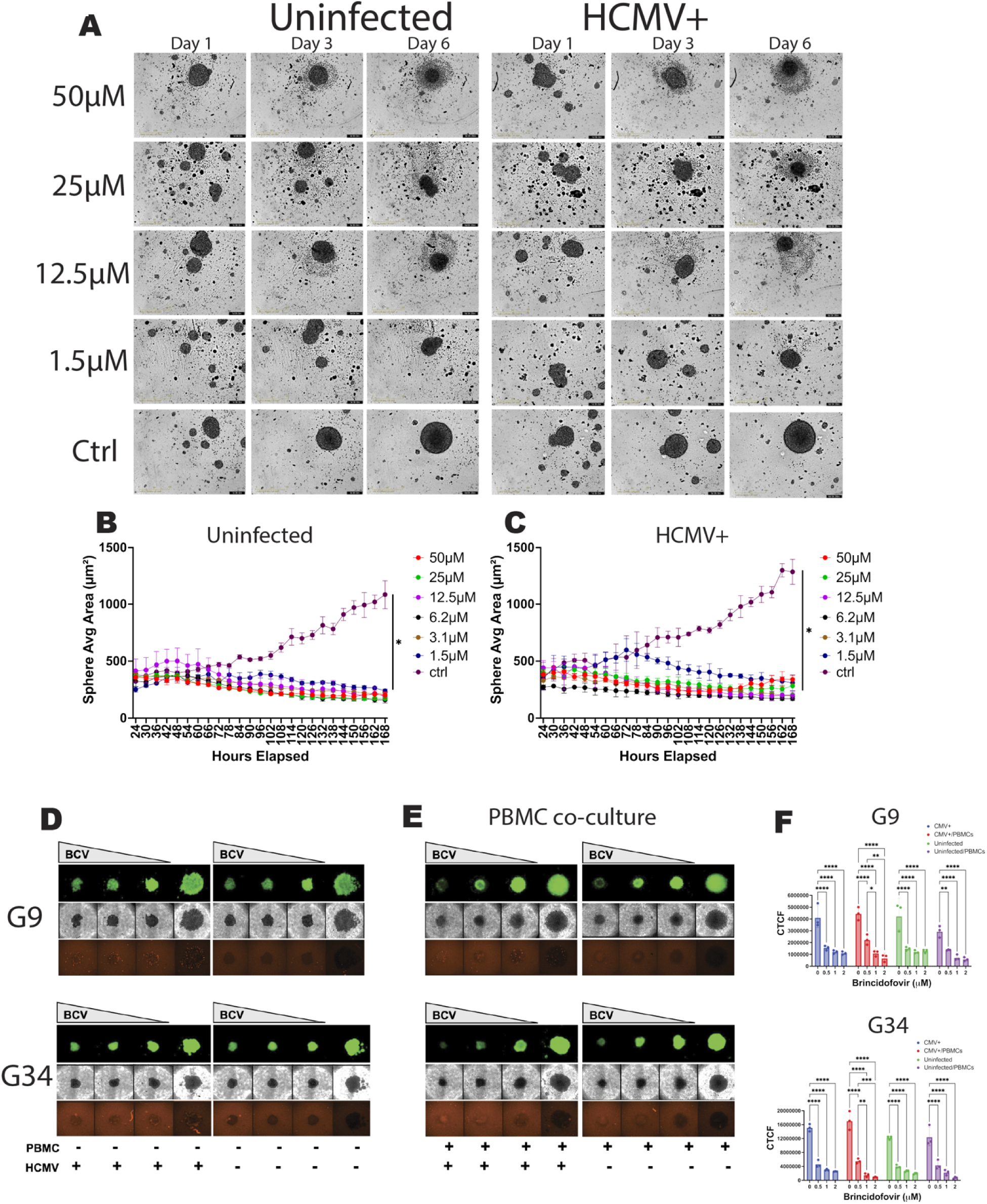
BCV prevents growth of patient derived neurospheres. (A) Representative HD phase contrast images of patient derived tumor neurospheres (G44) single spheroids, n=3 technical replicates per condition, in a 96-well ultra-low attachment round bottom plate comparing HCMV infected and uninfected controls with increasing concentrations of BCV. Images were acquired every 6 hrs using 10x objective. Scale bar: 400µm. (B) quantitation of sphere average area in µm^2^ of uninfected and (C) HCMV infected spheroids after treatment with BCV. (D) Representative images (4X) of single patient derived tumor neurospheres (G9-pCDH and G34-pCDH) on day 6 after HCMV infection and treatment with BCV at the following concentrations 2 µM, 1 µM, 0.5 µM and 0 µM. (E) Co-culture assays of patient derived GBM tumor neurospheres (G9-pCDH and G34-pCDH) after HCMV infection and co-culture with 20k healthy donor PBMCs. (F) Corrected total cell fluorescence (CTCF) of GFP+ tumor neurospheres. Statistical analysis was performed using two-way ANOVA followed by Tukey’s multiple comparisons test. Data are presented as mean ± SD. Statistical significance: * *p* < 0.05; p > 0.05, ns, not significant.

In order to understand the therapeutic activity of BCV we performed western blot analysis after CMV infection and BCV treatment in SB28 cells and patient derived tumor neurospheres (G44). We found a significant increase in phosphorylated H2AX (γ-H2AX) after BCV treatment in the three cell lines suggesting increased DNA damage driven by BCV (Figure 3C, D). This increase was observed to be dose dependent where BCV concentrations of 10µM, significantly increased the levels of γ-H2AX compared to 2µM in SB28 indicating an accumulation of DNA damage. Notably, in SB28 a compounding effect of BCV and CMV was observed with slightly higher γ-H2AX levels, however, this was not observed in the tumor neurospheres (Figure 3C, D). Treatment with increasing concentrations of BCV resulted in a clear dose-dependent increase in γ-H2AX foci formation in SB28 cells, as detected by immunofluorescence staining (Figure 3E).

Representative images display elevated numbers of discrete γ-H2AX -positive foci (red) within nuclide (DAPI, blue). Interestingly increased levels were observed in SB28 cells following MCMV infection, suggesting additive effects of viral infection and treatment-induced genotoxic stress (Figure 3F). Together, these findings are consistent with prior reports showing that CDV induces dose-dependent DNA damage in GBM cells^28^.

We next investigated whether BCV alters other cellular pathways in GBM. Interestingly the combination of CMV infection and BCV treatment increased cleaved PARP levels in SB28 cells and G44 neurospheres, indicating induction of apoptosis (Figure 3C, E). Given the role of c-Myc in promoting tumor cell proliferation and stemness, we further assessed its expression following treatment with BCV. BCV reduced c-Myc levels at higher concentration (10 µM) compared to lower dose (2 µM) in SB28 cells, consistent with decreased proliferative capacity (Figure 3C). In contrast, MCMV infection increased c-Myc expression in SB28 cells, whereas this effect was not observed in the G44 neurospheres, potentially reflecting differences in early viral infection kinetics. Together, these data suggest that BCV induces DNA damage-associated apoptosis signaling and suppresses proliferative pathways in GBM in a dose dependent manner.

### Brincidofovir prevents growth of patient-derived glioblastoma cells

BCV exerts direct molecular effects in GBM cells; therefore, to further assess its impact on tumor growth, we performed proliferation assays using an IncuCyte live cell imaging system. Patient-derived G44 GBM neurospheres were treated with increasing concentrations of BCV and monitored over 6 days (Figure 4A). We found that BCV inhibited the growth of GBM tumor neurospheres even at low concentrations (<1.5 µM) suggesting potent direct antitumor effects on GBM cell proliferation. The average sphere size did not change in the treated wells compared to the controls where a steady increase in sphere size was observed (Figure 4B) (*p* < 0.05). Interestingly, in the CMV infected tumor spheres we observed larger spheres compared to the uninfected control and at low doses of BCV (1.5 µM) the sphere size initially increased then after 72 hours decreased in size (Figure 4C) (*p* < 0.05). To confirm that BCV was killing these cells we performed cell viability assays and observed that BCV significantly reduced the viability of these tumor neurospheres (supplemental figure 2) (*p* < 0.05). Interestingly, we observed that HCMV increases the viability of these cells which was observed in both viability and proliferation assays (Figure 4C and Supplementary Figure 2). This viability increase was retained after treatment with TMZ and radiotherapy (Supplemental Figure 2) (*p* < 0.05). After treatment with BCV a reversal of this pro-growth phenotype was observed (Supplemental Figure 2). These data show that BCV impairs growth and viability of GBM patient-derived tumor cells.

We next evaluated whether BCV also affects additional patient-derived GBM neurospheres (G9-pCDH and G34-pCDH) and found that it similarly inhibited their growth (Figure 4D, E). Notably, CMV infection increased CTCF in G34-pCDH neurospheres, consistent with enhanced growth, whereas this effect was not observed in G9-pCDH cells (Figure 4F, G) (*p* < 0.05). We next assessed whether BCV enhances tumor cell killing in the presence of healthy donor PBMCs. PBMC co-culture altered neurosphere morphology, leading to a more compact, rounded phenotype without significant changes in size (Figure 4E). In addition, BCV modestly decreased CTCF in PBMC-co-cultured of tumor neurospheres (Figure 4F, G) (*p* < 0.05). Together, these data show that BCV suppresses growth across heterogenous GBM neurospheres and maintains activity in the presence of immune cells.

### Cytomegalovirus upregulates pro-tumorigenic proteins which are targeted by Brincidofovir

To identify protein mediators of CMV-driven tumor promotion in GBM neurospheres and determine whether BCV directly counteracts these effects, we performed proteomic profiling following CMV infection and BCV treatment. We observed a greater number of upregulated proteins in HCMV-infected tumor neurospheres compared to BCV-treated (Figure 5AB). Proteins upregulated in HCMV-infected neurospheres were enriched for previously described pro-tumorigenic factors, including Rho GTPase activating protein 12 (ARHGAP12)^38^, SERPINE1^39^ and the E3 ubiquitin-protein ligase Praja-2 (PJA2)^40^, among others (Figure 5C). Notably, BCV treatment reversed the expression of these proteins, resulting in their significant downregulation (Figure 5C). These data indicate that CMV remodels the GBM proteome toward a pro-tumorigenic, while BCV counteracts this program by suppressing key viral-induced protein networks.

**Figure 5:**
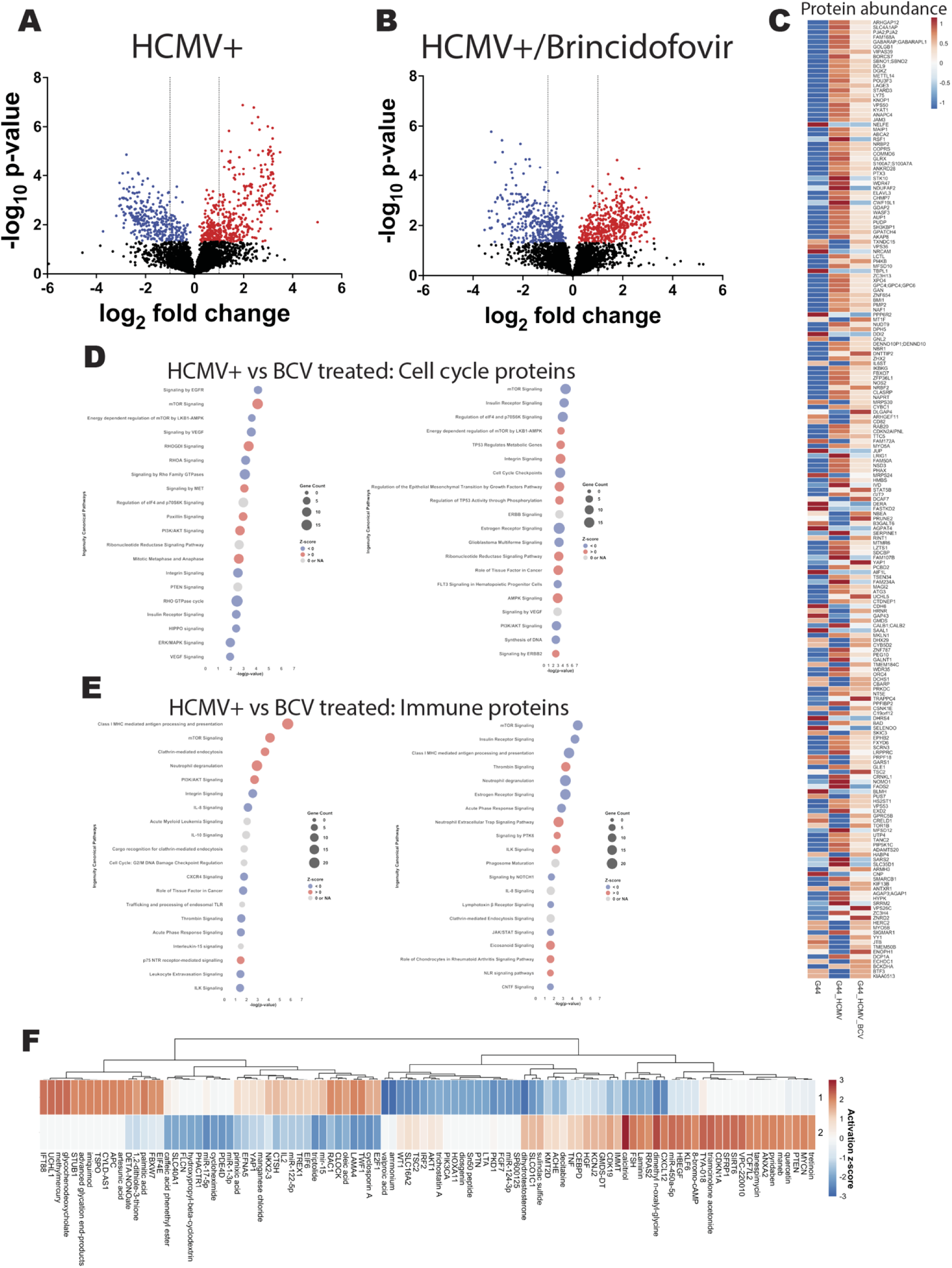
Proteomic analysis of HMCV infection and BCV treatment in patient derived tumor neurospheres G44. (A) volcano plot of differential protein abundance of uninfected and (B) HCMV infected with BCV treatment after 72hrs. Horizontal axis: log2 fold change; vertical axis -log10 (p-value). Points colored by significance (red=upregulated, blue=downregulated. (C) Heatmap of relative protein abundance across conditions (uninfected, HCMV infected, and BCV treated. Proteins shown in order of abundance with no clustering. Color scale: blue (low) red (high) relative to row mean. Generated using pheatmap in R. (D) Dot plot of top 20 enriched Ingenuity Canonical pathways of major cell cycle regulators and (E) immune regulators, ranked by -log10 (p-value) dot size: number of overlapping proteins; color=activation z-score (red=activated, z > 0; blue=inhibited, z < 0; gray=neutral/N/A). Generated using ggplot2 in R. (F) Horizontal clustered heatmap of upstream regulators by protein activation z-score comparing 1. HCMV infected to 2. BCV treated. Hierarchical clustering (complete linkage, Euclidean distance) was applied to proteins (column dendrogram; row (conditions) are not clustered. Color scale ranges (-3 blue, downregulated, 0 white, no change, and +3 red, upregulated. Generated using pheatmap in R.

**Figure 6:**
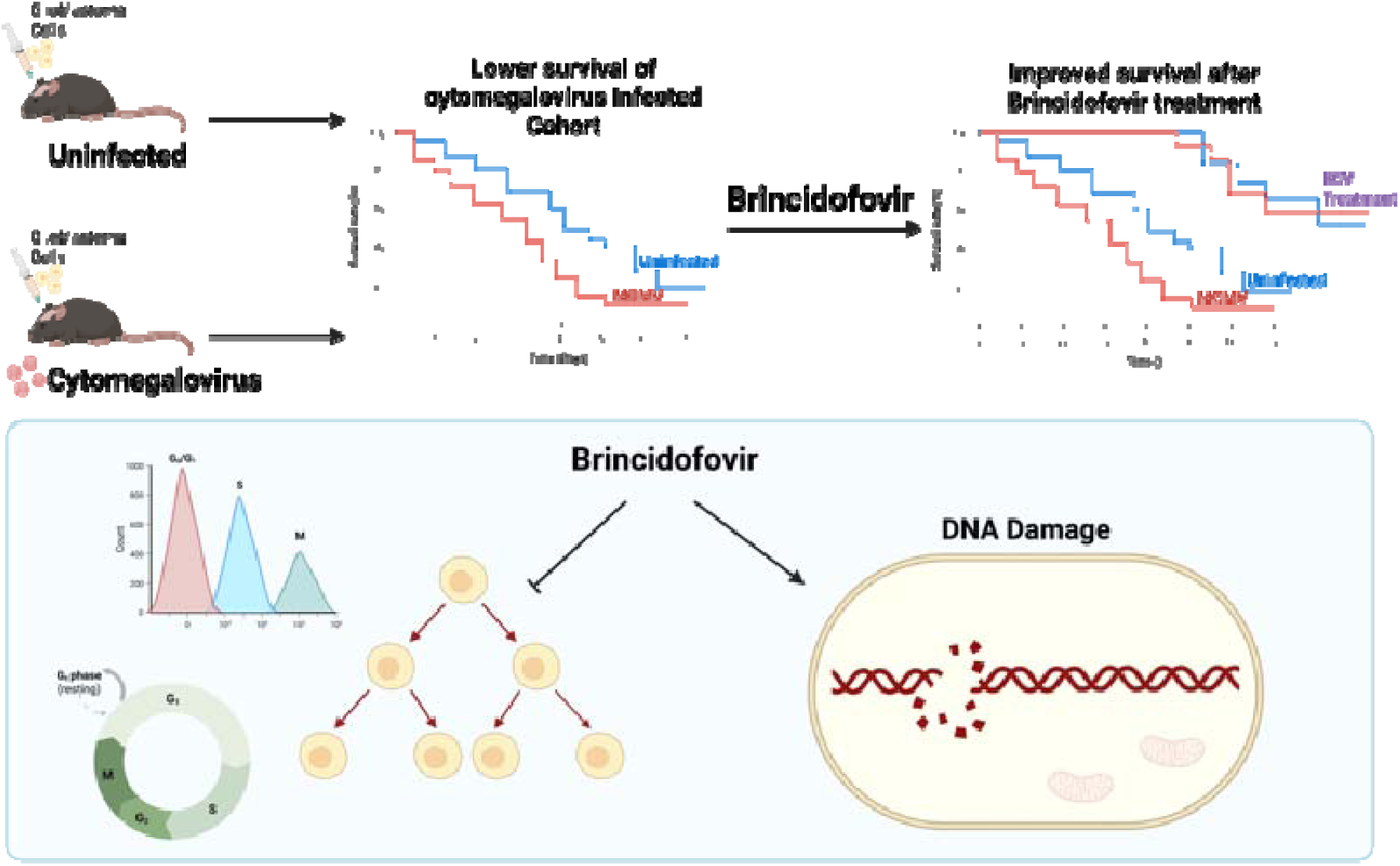
Graphical abstract

To characterize pathways altered by HCMV infection and BCV treatment, we performed Gene Ontology (GO) enrichment analysis comparing infected to BCV and treated tumor neurospheres. HCMV infection was associated with enrichment of proteins involved in mTOR and PI3K/AK signaling, as well as immune-related pathways including MHC class I antigen processing and presentation. These pro-tumorigenic pathways and immune-modulatory pathways were broadly downregulated following BCV treatment (Fig. 5D, E), indicating reversal of the HCMV-induced proteomic program. Upstream regulator analysis further predicted activation of tumor suppressors, including PTEN and CDKN1A, upon BCV treatment (Fig. 5F). Collectively, these data define a CMV-driven proteomic state in GBM that is counteracted by BCV, providing mechanistic insight into its antitumor activity.

## DISCUSSION

In this study, we report the promotion of GBM growth by murine cytomegalovirus infection in a clinically relevant murine model of GBM (SB28) which closely mimics human GBM. We show that CMV potentiates tumor growth resulting in significant clinical decline compared to naïve-uninfected controls. We demonstrate that treatment with the antiviral drug Brincidofovir prevents tumor growth and significantly improves long-term (<80 days) survival of tumor bearing mice when compared to untreated controls. Of note, mice treated with BCV did not exhibit onset clinical signs of decline due GBM tumor growth whereas all untreated mice developed clinical symptoms. Our data support previous findings of immune remodeling by CMV in other murine GBM models. We demonstrated that CMV infection promotes immune tumor inflammation in SB28 tumor bearing mice. In contrast, BCV reversed these immune changes and interestingly elicited B cell expansion, which may indicate increased long-term immune surveillance. Notably, there was a decrease in a subtype of Ly6C expressing macrophages after BCV treatment (cluster Fc-02) which indicates a reduction in tumor inflammation after treatment. Together, these data provide direct support of the importance of CMV in GBM tumor progression and to our knowledge this is the first report demonstrating the direct anti-tumor effects of Brincidofovir in a clinically relevant model of GBM. Currently only Valganciclovir has shown some clinical promise for GBM, therefore additional therapeutic drugs like BCV may be beneficial to test in the clinic.

In order to understand how BCV prevents tumor progression, we tested *in vitro* clinically relevant concentrations of BCV in various GBM lines. Similar to previous studies conducted using CDV, we found that BCV induces significant DNA damage and S-phase arrest in GBM. Interestingly, our data demonstrate that GBM cells infected with CMV then treated with BCV amplify the DNA damage. This suggests that GBMs with actively replicating virus may be more susceptible to BCV treatment. Our data support previous findings showing that incorporation of CDV into the DNA of proliferating tumor cells induces double-stranded DNA breaks and ultimately leads to tumor cell death^28^. Several reports have indicated that CMV is present in GBM; however, other studies have failed to detect CMV. Therefore, testing BCV in GBM patients may be feasible, as this FDA-approved drug could provide therapeutic benefit regardless of CMV status.

Here we provide evidence that BCV directly impacts GBM tumor cell growth through c-MYC, an important regulator of proliferation in GBM. At concentrations of 10 µM, c-Myc expression decreased in SB28 cells. These findings are supported by our observations of slower proliferation kinetics and increased killing in murine GBM cells (SB28) and tumor neurospheres (G44, G9-pCDH, G34-pCDH) following BCV treatment. Co-culture experiments using healthy donor PBMCs and tumor neurospheres showed that BCV inhibited tumor growth in the presence of peripheral immune cells. Notably, there was a trend towards increased neurosphere killing following the treatment with both PBMCs and BCV, although this did not reach statistical significance. These findings suggest that BCV may influence how immune cells target tumor cells, although additional studies are required to determine which immune cell populations are involved.

Previous reports suggest that cytomegalovirus induces changes at the molecular level of tumor cells^41–44^. Proteomic analysis of G44 tumor neurospheres following CMV infection revealed upregulation of several pro-tumorigenic proteins, including Rho GTPase activating protein 12 (ARHGAP12), which is involved in migration and invasion as well as other proteins that have been widely implicated in tumor progression of GBM, SERPINE1 and E3 ubiquitin-protein ligase Praja-2 (PJA2). After BCV treatment, these proteins were either significantly downregulated or no longer upregulated, suggesting that BCV targets several proteins upregulated following CMV infection. Pathway analysis revealed that HCMV upregulates several major signaling pathways, including mTOR, PI3K/AKT and MET^41^, which are subsequently suppressed following BCV treatment. Importantly, immune-related pathways altered by HCMV infection, which provide insight into the immune-inflamed tumor microenvironment, were also targeted by BCV treatment. These findings highlight the impact of active HCMV infection on tumor cell biology and identify molecular pathways that may be therapeutically targeted. These altered pathways provide evidence of HCMV oncomodulation and are supported by previous findings.

Although our findings provide new insights, there are limitations to this study. In particular, additional models are needed to determine whether CMV similarly impacts GBM growth across the heterogeneous spectrum of GBM. To address this limitation, we are preparing a manuscript describing additional models of CMV-driven tumorigenesis in GBM. Additional *in vivo* data is also necessary to determine if BCV similarly provides therapeutic benefit in other models of GBM.

In summary, our findings identify CMV as a biologically active driver of GBM progression and establish BCV as a dual-action therapeutic capable of targeting both viral and tumor-intrinsic mechanisms. These results support further clinical investigation of CMV-targeted strategies in GBM and suggest that viral oncomodulation represents a therapeutically actionable axis in brain tumors.

## MATERIALS AND METHODS

### Cell lines and virus stocks

SB28-Ohlfest cells were purchased from Leibniz Institute DSMZ-German Collection of Microorganisms and Cell Cultures (GmbH) (cat # ACC 880) and cultured in Dulbecco’s Modified Eagle Medium/F12 (Gibco) supplemented with 10% fetal bovine serum (Gibco) and 0.1% penicillin/streptomycin (Fisher). Patient-derived glioma stem cell-like tumor neurospheres G44, G9-pCDH, and G34-pCDH were obtained and cultured as previously described^45,46^, cells were maintained as neurospheres in complete Neurobasal medium (Gibco) supplemented with 2% B-

27 supplement (Fisher Scientific), 0.1% GlutaMax (Fisher Scientific), 20 ng/mL of human recombinant EGF (Peprotech), 20 ng/mL of human FGF (Peprotech) and 0.1% penicillin/streptomycin (Fisher). Tumor neurospheres were allowed to form overnight prior to experiments. Human cytomegalovirus TB40 strain expressing mCherry and eGFP were generated at Upstate Medical University. Viral BAC DNA isolation, transfection, expansion, and tittering were carried out as previously described (Carter et al., 2025). Briefly, BAC DNA was transfected into hTert MRC5-cells (10cm dish, ∼1 x 10e6 cells), allowed to reach 100% cytopathic effect, and then cell-associated and cell-free virus was isolated. One-tenth of the supernatant was then expanded on naïve NuFF-1 cells (9 15cm dishes, ∼4.5 x 10e7 cells). Cell-associated and cell-free virus was harvested by ultracentrifugation through a 20% sorbitol cushion. The viral pellet was then resuspended in full media containing 1.5% bovine serum albumin, flash frozen with liquid nitrogen, and stored at -80°C. Titers for each stock were then calculated using tissue culture infectious dose assay (TCID50). Viral growth kinetics were assessed in NuFF-1 cells by infecting at an MOI of 1 and collecting cell-free virus at 24, 48, 72, 96, and 120 hours post infection. Murine Cytomegalovirus (MCMV) lacking the m157 gene (Δm157)^47^ was provided by Ulrich Koszinowski (Ludwig-Maximilians-Universitat, Munich, Germany).

### In vivo studies

Pregnant C57BL/6 females (Envigo) were obtained at E11 for *in vivo* experiments. All procedures were performed in accordance with Institutional Animal Care and Use Committee (IACUC) guidelines (protocol 24-09-0004) and supported by the Center for Animal Resources and Education at Brown University. For survival studies, pups were infected at P2 with murine cytomegalovirus (MCMV) Δm157 (Smith strain) via intraperitoneal injection. At 10 weeks, infection was confirmed by serum IgG analysis. At 14 weeks of age, 1,500 SB28-Ohlfest cells in 3μL were implanted intracranially using stereotactic coordinates (2 mm right lateral, 1 mm frontal to bregma, and 3 mm deep). Cohorts included both male and female C57BL/6 mice. Seven days after implantation, mice received 20 mg/kg of Brincidofovir (Symbio) intraperitoneally twice weekly for a total of eight doses over one month, while control animals received vehicle phosphate buffered saline (PBS). Mice were housed in groups under identical conditions. Study endpoints included 20% weight loss and neurological symptoms assessed by clinical scoring.

### Proteomic analysis

Patient derived G44 tumor neurospheres were seeded at 2×10^6^ cells in technical duplicates an ultra-low adherent flask and allowed to form neurospheres for 24 hours then they were infected with 1 multiplicity of infection (MOI) of HCMV TB40 strain for 2 hours. Cells were then treated with 2 µM of Brincidofovir then left to incubated at 37 C 5% CO2 for 72 hours. Cell lysates were harvested in Ripa buffer (Fisher) containing 1x Protease Phosphatase Inhibitor (Fisher) then stored at -80 C. For proteomic analysis,100 μg of protein was subjected to S-Trap Micro Digestion and Clean-up (Protifi, C02-micro-40). Approximately 20 μL of Lysis Buffer (10% SDS, Invitrogen) in 50 mM Triethylammonium Bicarbonate Buffer pH 7.5 (\Sigma) was added to each sample. Proteins were reduced using Dithiothreitol (Sigma) and incubated at 55°C for 45 minutes with mixing. Samples were alkylated using 20 mM Iodoacetamide (Fisher) and incubated at room temperature in the dark for 30 minutes then acidified using Phosphoric Acid. 6 volumes of ice-cold Binding Buffer were added to each sample before the entire sample was transferred to an S-Trap spin column and centrifuged for 30 seconds at 4000xg. Samples were washed with binding buffer then Trypsin (Promega) was added to each micro spin column. Samples were moved to a humidified chamber in a 37°C incubator and left overnight (18 hours). After incubation samples were left to cool then 50 μL of 50 mM TEAB was added. Hydrophilic peptides were eluted with 50 μL of LC/MS grade Water (Honeywell) and 0.1% Formic Acid (Fisher). Hydrophobic peptides were eluded in 50 μL of 50% LC/MS Grade Acetonitrile (Fisher). Each sample was concentrated using a speed vacuum concentrator (Fisher) for approximately 90 minutes. With solvent removed, samples were reconstituted in 50μL of Solvent A (Water with 0.1% Formic Acid) and spiked with indexed retention time peptides (Biognosys) in a 1:50 dilution to monitor UHPLC performance.

Reconstituted Peptide samples were then injected in duplicate onto a QExactive Orbitrap LC/MS system (Fisher) and separated over a 100-minute gradient consisting of water + 0.1% Formic Acid (Solvent A), and Acetonitrile + 0.1% Formic Acid (Solvent B) using a 300 μm x 5 mm C18 trap-elute set-up (Fisher) on an Ultimate 3000 RSLC UHPLC system. A 25cmx75 μm ID EasySpray capillary analytical column packed with C18 2 μm resin (Fisher) was used to separate digested peptides. The method consisted of 2% B (loading pump flow rate 6 μL/min, nano pump flow rate 300nl/min) for 5 min to load the trap column and elution using 2%B for 5 mins (300 nl/min), 2-20%B for 45mins (200 nl/min), and 20-40% over 50 mins(200 nl/min). The mass spectrometer acquisition utilized a Full MS/ddMS2 (Top20) centroid experiment. Full MS parameters used a default charge state of 2, 1 microscan, resolution of 70,000, AGC target of 1×10^6^, 30ms Maximum Injection time, in a scan range of 400-1800m/z. ddMS2 parameters were 1 microscan, resolution of 17,500, AGC target of 5×10^4^, a max injection time of 75ms, a loop count of 20, Top 20 precursors, Isolation window of 2.0 m/z, normalized collision energy of 25. Data dependent settings consisted of a minimum AGC target of 5×10^2^, 6.7×10^3^ intensity threshold, unassigned, 1,6-8, and >8 charge exclusion, preferred peptide match, isotope exclusion, 30 seconds of dynamic exclusion.

*Label Free Quantitation and SpectroMine Search Parameters,* .raw files were processed using Spectromine (Version 4.4.240326) using 1% peptide and protein FDR search constraints, all settings where system defaults against the reference Homo sapiens protein. fasta from UNIPROT (20230316, 42,442 entries, canonical with isoforms). Variable modifications consisted of oxidation (M), deamidation (N), and protein N-terminal Acetylation. Carbamidomethylation was the sole static modification for the Trypsin search. The exported protein group report was further processed in excel and relative abundance was calculated as the intensity of each protein divided by the sum of all intensities for each given sample.

*Missing value Imputation,* Random Forest (RF) imputation was performed as described in Jin et al^48^ using “missForest” in R package^49^. After imputation log2 fold change was calculated in excel along with Welch’s t-test for p-value analysis. Graphpad prism was using to generate volcano plots of all upregulated and downregulated proteins.

### Gene ontology (GO) analysis

Gene ontology analysis was performed using the Qiagen Ingenuity Pathway Analysis software. Excel file containing the data set generated above was imported into the software and enrichment analysis was performed. GO pathway plots for immune and cell cycle were plotted using the ggplot2 package in R and the heatmap of upstream regulators using the pheatmap package in R.

### *In vitro* cell proliferation assays

SB28-Olhfest were seeded at 1×10^3^ cells per well and G44 cells were seeded at 5×10^3^ in technical triplicates using complete medium then after 24 hours were infected with 1 MOI murine cytomegalovirus Δm157 smith strain. After 2 hours the cells were treated with increasing concentrations of Brincidofovir. Plates were kept in an Incucyte (Sartorius) for live cell imaging and analysis.

### Cell cycle analysis

For cell cycle analysis, 2×10^4^ SB28-ohlfest cells and 1×10^5^ patient derived G44 tumor neurospheres were plated in a six-well plate in complete medium. Cells were then infected with 1 MOI of HCMV TB40 or MCMV Δm157 for 2 hours. After infection cells were treated with 2µM of Brincidofovir and incubated for 48 hours at 37 C 5% CO2. Next, cells were collected and washed three times with PBS and stained with propidium iodide (1 mg/ml) at 1:1000 dilution for 30 min in the dark. Cells were taken for analysis in a BD Fortessa cytometer, and data were analyzed using a FlowJo software (BD Biosciences).

### Immunoblot assays

Patient derived tumor neurospheres (G44) were seeded at 2×10^6^ then incubated for 24 hours to allow for sphere formation. Cells were then infected with 1 MOI of HCMV TB40 for 2 hours then treated with 2 µM of Brincidofovir. Similarly, SB28-ohlfest cells were seeded at 1×10^5^ cells for 24hours then infected with murine cytomegalovirus (MCMV) at 1 MOI followed by treatment with BCV (2 µM). Cell lysates were collected 72 hours post treatment in Ripa buffer (Fisher scientific) containing 1x Protease Phosphatase Inhibitor (Fisher) then stored at -80 C. Total protein concentration was measured using a Pierce BCA Protein Assay Kit (Fisher) according to manufacturer’s instructions. A Molecular Devices SpectraMax M2 plate reader was used at 660nm absorbance to measure protein concentrations. For each sample 10 µg of protein was incubated with 1× Laemmli sample buffer (Bio-Rad) at 95°C for 5 min before loading onto 10% Mini-PROTEAN TGX precast protein gel (Bio-Rad). A PageRuler Plus Pre-stained Protein Ladder (Fisher) was used as a ladder. Blocking was performed in 5% milk with 0.1% Triton X-100 in TBST (Gibco) for 1 hour at room temperature under shaking. Primary antibodies used were incubated in the cold under shaking overnight. Primary antibodies used were against γ-H2AX (Cell Signaling Technologies 1:1000), PARP/cleaved PARP (Cell Signaling 1:1000), c-MYC (cell signaling 1:1000), and β-actin (Cell Signaling Technology 1:1000). GelAnalyzer 23.1.1 software was used for Western blot quantification. For each blot, the chemiluminescence image was analyzed. The setting for dark spots on a light background was selected. Lanes were detected using the equal width option, and the region of interest containing the protein was defined for lane detection. The following settings were applied: minimal height: 5, minimal slope (degrees): 29, slope break in threshold (degrees), and profile smoothing radius: 1. Baseline detection used the rolling ball method, with a peak width tolerance of 10% of the profile length.

Bands and baselines were adjusted to capture the entire peak. Normalization was performed by dividing the peak value of the protein of interest to the housekeeping protein.

### Immunofluorescence staining

Mice were euthanized using CO_2_ inhalation then brains were collected and immediately fixed in 10% formalin for 72 hours on a rotator at 4 C. The brains were transferred to 30% sucrose and sodium azide then stored at 4 C. Prior to cryosectioning brains were frozen at -80 C for more than 30 minutes, embedded in Optimal Cutting Temperature (OCT) compound (Fisher) and transferred to -27 C in a cryostat (Leica CM1950). All mouse brain slides were obtained from 20μm frozen sections. Brain tissue sections were permeabilized using 0.01% Triton X100 (Fisher) and blocked for 1hr using normal donkey serum (Fisher). After incubation primary antibodies were added and incubated overnight at room temperature. For primary antibody detection, species-matched fluorophore-coupled antibodies were incubated for 1 hour at room temperature. Slides were then covered with xylene mounting medium (Fisher) and a cover slip was placed over the slide. High-resolution confocal fluorescent microscopy was performed using a Zeiss LSM 710 confocal microscope system and visualized using ZEN Zeiss Imaging software and Olympus VS200 whole slide scanner. The following antibodies were used: mouse anti-CD31 (Cell Signaling Technologies, catalog number 3528S, 1:50), anti-m123 (clone CROMA101) (Center for Proteomics University of Rijeka, catalog number HR-mCMV-08, 1:50).

### Leucocyte co-culture assays

PBMCs were obtained from healthy human donors as approved by the Institutional Review Board (IRB) at Brigham and Women’s Hospital (all samples were de-identified) and isolated with Ficoll-Paque PLUS density gradient medium (GE Healthcare Life Sciences) according to the manufacturer’s instructions. A single-cell suspension of patient derived GBM cells (G9-pCDH and G34-pCDH) were seeded at 5×10^3^ cells/well in ultra-low-attachment 96-well plates (Corning) and incubated for 24 hours to allow sphere formation. Spheres were then infected with HCMV TB40 for 2 hours. Following infection PBMCs were then added with increasing Brincidofovir concentrations. Cells were then incubated for 6 days of co-culture. Microscope images of the spheres were taken daily (ImageXpress Micro Confocal High-content Imaging System), and the spheres’ fluorescence was measured in ImageJ.

### Murine tumor-infiltrating leukocyte Isolation

The right hemisphere bearing the tumor was collected from each mouse at end points. N=3/4 mice per cohort were included in the analysis. A tumor dissociation kit for mouse (Miltenyi Biotec) was used for isolation of tumor infiltrating leukocytes according to the manufacturer’s instructions. Harvested leukocytes were stored at −80°C in Bambanker prior to downstream analysis.

### Mass cytometry (CyTOF)

Samples were processed as previously described briefly, samples were thawed at 37°C with RMPI medium supplemented with fetal bovine serum, Glutamax, and antibiotic-antimycotic. Then MEM was added with HEPES, (4-(2-hydroxyethyl)-1-piperazineethanesulfonic acid, 2-mercaptoethanol, sodium heparin, and benzonase nuclease. Samples were then mixed with PBS and 0.5 and 2.0×10^6^ cells were analyzed. Samples were fixed with 0.2% formaldehyde before staining. Mouse anti-CD16/32 antibody Fc-receptor blocking reagent (BioLegend) was used in cell staining buffer. All antibodies were obtained from the Harvard Medical Area CyTOF Antibody Resource and Core (Boston, Massachusetts, USA).

### Quantitation and statistical analysis

Graphs and statistical analysis were performed using GraphPad Prism (Prism 5) or Microsoft Excel. Figure legends show details of statistical tests, including numbers and p values.

Independent experiments were performed to demonstrate reproducibly of results. A value of *p* ≤ 0.05 was considered statistically significant (*p* ≤ 0.05 *, p < 0.01 **, p < 0.001 ***).

## Acknowledgements

We thank Nicholas DaSilva (Proteomics Core Facility), Christoph Schorl (Genomics Core Facility) for generous advice, assistance, and reagents.

## Author Contributions

NBM and SEL designed the study. NBM performed all experiments, analyzed, and interpreted the data. NBM wrote the original manuscript. All co-authors provided extensive revisions. PV, WH, AS, JC, MS assisted with mouse surgeries. JP assisted with western blot experiments. AJ and AA assisted with cell cycle experiments and immunofluorescence imaging. MV and JAL assisted with CyTOF experiments. EP provided murine cytomegalovirus. EAM provided human cytomegalovirus. MH provided Brincidofovir. SEL and CHC acquired funding.

## Data and Materials Availability

All data are available in the manuscript or the supplementary material. Correspondence and requests for materials should be addressed to SEL

## References

1 Louis, D. N. et al. The 2021 WHO Classification of Tumors of the Central Nervous System: a summary. Neuro Oncol 23, 1231–1251 (2021). 10.1093/neuonc/noab106

2 De Silva, M. I., Stringer, B. W. & Bardy, C. Neuronal and tumourigenic boundaries of glioblastoma plasticity. Trends Cancer 9, 223–236 (2023). 10.1016/j.trecan.2022.10.010

3 Alexander, B. M. & Cloughesy, T. F. Adult Glioblastoma. J Clin Oncol 35, 2402–2409 (2017). 10.1200/JCO.2017.73.0119

4 Stupp, R. et al. Radiotherapy plus concomitant and adjuvant temozolomide for glioblastoma. N Engl J Med 352, 987–996 (2005). 10.1056/NEJMoa043330

5 Zhao, W. et al. Exploring tumor-associated macrophages in glioblastoma: from diversity to therapy. NPJ Precis Oncol 9, 126 (2025). 10.1038/s41698-025-00920-x

6 Gonzalez-Tablas Pimenta, M., et al. Tumor cell and immune cell profiles in primary human glioblastoma: Impact on patient outcome. Brain Pathol 31, 365–380 (2021). 10.1111/bpa.12927

7 Dasari, V., Smith, C., Schuessler, A., Zhong, J. & Khanna, R. Induction of innate immune signatures following polyepitope protein-glycoprotein B-TLR4&9 agonist immunization generates multifunctional CMV-specific cellular and humoral immunity. Hum Vaccin Immunother 10, 1064–1077 (2014). 10.4161/hv.27675

8 Nair, S. K. et al. Recognition and killing of autologous, primary glioblastoma tumor cells by human cytomegalovirus pp65-specific cytotoxic T cells. Clin Cancer Res 20, 2684–2694 (2014). 10.1158/1078-0432.CCR-13-3268

9 Scheer, I. et al. Prospective Evaluation of CD45RA+/CCR7- Effector Memory T (T(EMRA)) Cell Subsets in Patients with Primary and Secondary Brain Tumors during Radiotherapy of the Brain within the Scope of the Prospective Glio-CMV-01 Clinical Trial. Cells 12 (2023). 10.3390/cells12040516

10 Griffiths, P. & Reeves, M. Pathogenesis of human cytomegalovirus in the immunocompromised host. Nat Rev Microbiol 19, 759–773 (2021). 10.1038/s41579-021-00582-z

11 Dupont, L. & Reeves, M. B. Cytomegalovirus latency and reactivation: recent insights into an age old problem. Rev Med Virol 26, 75–89 (2016). 10.1002/rmv.1862

12 Jarvis, M. A. & Nelson, J. A. Human cytomegalovirus persistence and latency in endothelial cells and macrophages. Curr Opin Microbiol 5, 403–407 (2002). 10.1016/s1369-5274(02)00334-x

13 Cobbs, C. S. et al. Human cytomegalovirus infection and expression in human malignant glioma. Cancer Res 62, 3347–3350 (2002).

14 Dziurzynski, K. et al. Glioma-associated cytomegalovirus mediates subversion of the monocyte lineage to a tumor propagating phenotype. Clin Cancer Res 17, 4642–4649 (2011). 10.1158/1078-0432.CCR-11-0414

15 Rahbar, A. et al. Human cytomegalovirus infection levels in glioblastoma multiforme are of prognostic value for survival. J Clin Virol 57, 36–42 (2013). 10.1016/j.jcv.2012.12.018

16 Lucas, K. G., Bao, L., Bruggeman, R., Dunham, K. & Specht, C. The detection of CMV pp65 and IE1 in glioblastoma multiforme. J Neurooncol 103, 231–238 (2011). 10.1007/s11060-010-0383-6

17 Foster, H. et al. Human cytomegalovirus seropositivity is associated with decreased survival in glioblastoma patients. Neurooncol Adv 1, vdz020 (2019). 10.1093/noajnl/vdz020

18 Mercado, N. B. et al. Clinical implications of cytomegalovirus in glioblastoma progression and therapy. NPJ Precis Oncol 8, 213 (2024). 10.1038/s41698-024-00709-4

19 Fornara, O. et al. Cytomegalovirus infection induces a stem cell phenotype in human primary glioblastoma cells: prognostic significance and biological impact. Cell Death Differ 23, 261–269 (2016). 10.1038/cdd.2015.91

20 Batich, K. A. et al. Long-term Survival in Glioblastoma with Cytomegalovirus pp65-Targeted Vaccination. Clin Cancer Res 23, 1898–1909 (2017). 10.1158/1078-0432.CCR-16-2057

21 Nair, S. K., Sampson, J. H. & Mitchell, D. A. Immunological targeting of cytomegalovirus for glioblastoma therapy. Oncoimmunology 3, e29289 (2014). 10.4161/onci.29289

22 Prins, R. M., Cloughesy, T. F. & Liau, L. M. Cytomegalovirus immunity after vaccination with autologous glioblastoma lysate. N Engl J Med 359, 539–541 (2008). 10.1056/NEJMc0804818

23 Stragliotto, G., Pantalone, M. R., Rahbar, A., Bartek, J. & Soderberg-Naucler, C. Valganciclovir as Add-on to Standard Therapy in Glioblastoma Patients. Clin Cancer Res 26, 4031–4039 (2020). 10.1158/1078-0432.CCR-20-0369

24 Krenzlin, H. et al. Cytomegalovirus promotes murine glioblastoma growth via pericyte recruitment and angiogenesis. J Clin Invest 129, 1671–1683 (2019). 10.1172/JCI123375

25 Genoud, V. et al. Responsiveness to anti-PD-1 and anti-CTLA-4 immune checkpoint blockade in SB28 and GL261 mouse glioma models. Oncoimmunology 7, e1501137 (2018). 10.1080/2162402X.2018.1501137

26 Montoya, M. et al. Interferon regulatory factor 8-driven reprogramming of the immune microenvironment enhances antitumor adaptive immunity and reduces immunosuppression in murine glioblastoma. Neuro Oncol 26, 2272–2287 (2024). 10.1093/neuonc/noae149

27 Hatae, R. et al. Enhancing CAR-T cell metabolism to overcome hypoxic conditions in the brain tumor microenvironment. JCI Insight 9 (2024). 10.1172/jci.insight.177141

28 Hadaczek, P. et al. Cidofovir: a novel antitumor agent for glioblastoma. Clin Cancer Res 19, 6473–6483 (2013). 10.1158/1078-0432.CCR-13-1121

29. 29 in Trends in Antiviral Drug Development 447–500 (2025).

30 Ho, E. S., Lin, D. C., Mendel, D. B. & Cihlar, T. Cytotoxicity of antiviral nucleotides adefovir and cidofovir is induced by the expression of human renal organic anion transporter 1. J Am Soc Nephrol 11, 383–393 (2000). 10.1681/ASN.V113383

31 Florescu, D. F. & Keck, M. A. Development of CMX001 (Brincidofovir) for the treatment of serious diseases or conditions caused by dsDNA viruses. Expert Rev Anti Infect Ther 12, 1171–1178 (2014). 10.1586/14787210.2014.948847

32 Donaldson, A. et al. Broad-spectrum antiviral brincidofovir inhibits Epstein-Barr virus and related gammaherpesvirus in human and nonhuman primate cells. J Clin Invest 136 (2026). 10.1172/JCI195764

33 Painter, W. et al. First pharmacokinetic and safety study in humans of the novel lipid antiviral conjugate CMX001, a broad-spectrum oral drug active against double-stranded DNA viruses. Antimicrob Agents Chemother 56, 2726–2734 (2012). 10.1128/AAC.05983-11

34 Marty, F. M. et al. CMX001 to prevent cytomegalovirus disease in hematopoietic-cell transplantation. N Engl J Med 369, 1227–1236 (2013). 10.1056/NEJMoa1303688

35 Toth, K. et al. Hexadecyloxypropyl-cidofovir, CMX001, prevents adenovirus-induced mortality in a permissive, immunosuppressed animal model. Proc Natl Acad Sci U S A 105, 7293–7297 (2008). 10.1073/pnas.0800200105

36 Marcelin, J. R., Beam, E. & Razonable, R. R. Cytomegalovirus infection in liver transplant recipients: updates on clinical management. World J Gastroenterol 20, 10658–10667 (2014). 10.3748/wjg.v20.i31.10658

37 Sanford, D. C. et al. Pivotal animal efficacy studies supporting brincidofovir licensure under the FDA animal rule. Antiviral Research 244, 106302 (2025). 10.1016/j.antiviral.2025.106302

38 Cheng, V. W. T. et al. ARHGAP12 and ARHGAP29 exert distinct regulatory effects on switching between two cell morphological states through GSK-3 activity. Cell Rep 44, 115361 (2025). 10.1016/j.celrep.2025.115361

39 Seker, F. et al. Identification of SERPINE1 as a Regulator of Glioblastoma Cell Dispersal with Transcriptome Profiling. Cancers (Basel) 11 (2019). 10.3390/cancers11111651

40 Lignitto, L. et al. Proteolysis of MOB1 by the ubiquitin ligase praja2 attenuates Hippo signalling and supports glioblastoma growth. Nat Commun 4, 1822 (2013). 10.1038/ncomms2791

41 Krenzlin, H. et al. Cytomegalovirus infection of glioblastoma cells leads to NF-kappaB dependent upregulation of the c-MET oncogenic tyrosine kinase. Cancer Lett 513, 26–35 (2021). 10.1016/j.canlet.2021.05.005

42 van Senten, J. R. et al. The human cytomegalovirus-encoded G protein-coupled receptor UL33 exhibits oncomodulatory properties. J Biol Chem 294, 16297–16308 (2019). 10.1074/jbc.RA119.007796

43 Kumar, A. et al. The Human Cytomegalovirus Strain DB Activates Oncogenic Pathways in Mammary Epithelial Cells. EBioMedicine 30, 167–183 (2018). 10.1016/j.ebiom.2018.03.015

44 Heukers, R. et al. The constitutive activity of the virally encoded chemokine receptor US28 accelerates glioblastoma growth. Oncogene 37, 4110–4121 (2018). 10.1038/s41388-018-0255-7

45 Jimenez-Macias, J. L. et al. Modulation of blood-tumor barrier transcriptional programs improves intratumoral drug delivery and potentiates chemotherapy in GBM. Sci Adv 11, eadr1481 (2025). 10.1126/sciadv.adr1481

46 Zdioruk, M. et al. PPRX-1701, a nanoparticle formulation of 6’-bromoindirubin acetoxime, improves delivery and shows efficacy in preclinical GBM models. Cell Rep Med 4, 101019 (2023). 10.1016/j.xcrm.2023.101019

47 Price, R. L. et al. Cytomegalovirus contributes to glioblastoma in the context of tumor suppressor mutations. Cancer Res 73, 3441–3450 (2013). 10.1158/0008-5472.CAN-12-3846

48 Jin, L. et al. A comparative study of evaluating missing value imputation methods in label-free proteomics. Sci Rep 11, 1760 (2021). 10.1038/s41598-021-81279-4

49 Stekhoven, D. J. & Bühlmann, P. MissForest--non-parametric missing value imputation for mixed-type data. Bioinformatics 28, 112–118 (2012). 10.1093/bioinformatics/btr597

