## Supplemental Figures for "Preclinical evaluation of Brincidofovir in glioblastoma demonstrates improved long term-survival and cytomegalovirus-dependent and independent effects"

Supplemental Figure 1: **Brincidofovir extends survival of SB28 tumor bearing mice. (A)**Representative in vivo bioluminescence imaging (BLI) of SB28-Ohlfest tumor bearing mice. SB28 cells stably expressing firefly luciferase were stereotactically implanted into the right striatum. Imaging was performed 2 weeks post tumor implantation for vehicle control top and BCV treated bottom. Mice were imaged 10 minutes post intraperitoneal D-luciferin (150mg/kg) administration. Color scale indicates photons/s/cm2; increasing signal reflects tumor growth. **(B)** Kaplan-Meier survival curve of vehicle control (n=7) and BCV treated (n=5) tumor bearing mice. $p < 0.0057$, log-rank test.

Supplemental figure 2**: Brincidofovir impacts viability of SB28 Cells. (A)** SB28 cell viability compared to GL261 after treatment with increasing concentrations of BCV and luciferase plate readout for viability. **(B)** SB28 and GL261 cell viability after MCMV infection and treatment with increasing concentrations of BCV. Below is the luciferase plate readout for viability. Images were acquired after 72hrs using IVIS. n=3 technical replicates per condition.

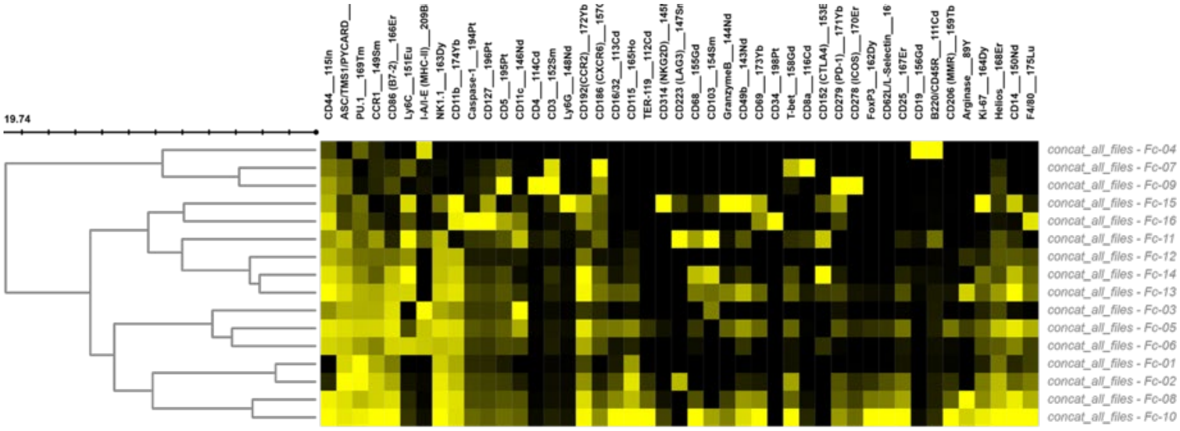


Supplemental figure 3: Heatmap of CyTOF marker expression across FlowSOM-derived clusters in four cohorts of tumor bearing mice 1. naïve 2. MCMV infected 3. Naïve BCV treated 4. MCMV infected BCV treated.

Supplemental figure 4: **Proteomic analysis of G44 tumor neurospheres reveals targeted HCMV proteins by BCV**. **(A)** Heatmap of HCMV protein abundance in 1. Uninfected 2. HCMV+ and 3. HCMV+/BCV treated tumor neurospheres. (B) Table describing HCMV genes and their function found in G44 tumor neurospheres after HCMV infection.
